# Genetic contributions to self-reported tiredness

**DOI:** 10.1101/047290

**Authors:** Vincent Deary, Saskia P Hagenaars, Sarah E Harris, W David Hill, Gail Davies, David CM Liewald, International Consortium for Blood Pressure GWAS, CHARGE consortium Aging and Longevity Group, Andrew M McIntosh, Catharine R Gale, Ian J Deary

## Abstract

Self-reported tiredness and low energy, often called fatigue, is associated with poorer physical and mental health. Twin studies have indicated that this has a heritability between 6% and 50%. In the UK Biobank sample (N = 108 976) we carried out a genome-wide association study of responses to the question, “Over the last two weeks, how often have you felt tired or had little energy?” Univariate GCTA-GREML found that the proportion of variance explained by all common SNPs for this tiredness question was 8.4% (SE = 0.6%). GWAS identified one genome-wide significant hit (Affymetrix id 1:64178756_C_T; p = 1.36 x 10-11). LD score regression and polygenic profile analysis were used to test for pleiotropy between tiredness and up to 28 physical and mental health traits from GWAS consortia. Significant genetic correlations were identified between tiredness and BMI, HDL cholesterol, forced expiratory volume, grip strength, HbA1c, longevity, obesity, self-rated health, smoking status, triglycerides, type 2 diabetes, waist-hip ratio, ADHD, bipolar disorder, major depressive disorder, neuroticism, schizophrenia, and verbal-numerical reasoning (absolute rg effect sizes between 0.11 and 0.78). Significant associations were identified between tiredness phenotypic scores and polygenic profile scores for BMI, HDL cholesterol, LDL cholesterol, coronary artery disease, HbA1c, height, obesity, smoking status, triglycerides, type 2 diabetes, and waist-hip ratio, childhood cognitive ability, neuroticism, bipolar disorder, major depressive disorder, and schizophrenia (standardised β’s between −0.016 and 0.03). These results suggest that tiredness is a partly-heritable, heterogeneous and complex phenomenon that is phenotypically and genetically associated with affective, cognitive, personality, and physiological processes.

“Hech, sirs! But I’m wabbit, I’m back frae the toon;

I ha’ena dune pechin’—jist let me sit doon.

From Glesca’

By William Dixon Cocker (1882-1970)

## Introduction

The present study examines genetic contributions to how the UK Biobank’s participants answered the question, “Over the last two weeks, how often have you felt tired or had little energy?” Ideal questionnaire items don’t have conjunctions, but the “or” is understandable here, and it may even allow capture of both peripheral and central fatigue. The first and last authors of the present study grew up in South Lanarkshire in Scotland, where fatigue was often self-reported in terms of feeling “wabbit”. The Scots word wabbit encompasses both peripheral fatigue, the muscle weakness after a long walk, and central fatigue, the reduced ability to initiate and/or sustain mental and physical activity, such as we might experience whilst having flu. Throughout the paper we refer mainly to the single English words “fatigue” and/or “tiredness” as the construct captured by the question, but the Scottish vernacular word is a good reminder that these constructs are multi-dimensional.

Fatigue is a common complaint. In a Dutch adult general population survey with 9375 respondents, 4.9% reported short term fatigue (<6 months duration), 30.5% chronic fatigue (> 6 months duration), and 1% fulfilled diagnostic criteria for Chronic Fatigue Syndrome (CFS) ^1^. These findings are similar to a London-based survey of general practice patients in England, aged 18-45, with 15283 respondents, where 36.7% reported substantial fatigue, 18.3% substantial fatigue of > 6 months duration and 1% fulfilled the diagnostic criteria for CFS ^2^. Two other large surveys of US workers and community dwelling adults aged 51 and over report fatigue rates of 37.9% (two week period prevalence) and 31.2% (one week period prevalence) respectively ^3, 4^. In an early review of fatigue epidemiology, Lewis and Wessely^5^ argue that fatigue “is best viewed on a continuum”, and the continuous distribution of fatigue in the general population is supported by the Pawlikowska et al. study^2^. Fatigue is also a common presentation in primary care. In a survey of 1428 consultations to 89 GPs in Ireland, fatigue prevalence was 25% and the main reason for attendance in 6.5% ^6^.

Demographically, higher levels of self-reported fatigue are associated with female gender, lower socio-economic status^1^ and poorer self-rated health status ^7^. There are less clear associations between age and fatigue, with some studies reporting a small but significant positive correlation between age and fatigue ^2^, whereas others report no association ^6^ or a negative association ^1, 7^.There is a clearer link with the Fried phenotype of frailty in older adults^8^, which has significant associations with mortality. The frailty phenotype comprises weakness (as measured by grip strength), weight loss, reduced mobility, reduced walking speed and fatigue^9^.

Fatigue is associated with a number of lifestyle-related factors and conditions. Smoking is a risk factor for fatigue^10^ and fatigue has strong cross-sectional associations with Type 2 Diabetes^11^ and increased BMI^12^. Fatigue is consistently associated with poorer physical and mental health status. It is one of the most common symptom complaints of cancer patients. For those undergoing treatment, prevalence estimates vary between 25% and 99%, and 25 to 30% of survivors report long term fatigue^13^. Fatigue is also a significant symptom of, to name just a few conditions, primary biliary cirrhosis (PBC)^14^, multiple sclerosis^15^, rheumatoid arthritis^16^, primary Sjogren’s syndrome^17^, and Parkinson’s disease^18^. It is associated with chronic disease in general, and there is a linear relationship between number of chronic diseases and self-reported fatigue^19^. Fatigue is also strongly associated with depression^20^, with self-reported psychological distress^2, 21^, and with the personality trait of neuroticism^22, 23^.

Research into the biological mechanisms of fatigue has focussed on a few key areas. Fatigue is associated with the cytokine-mediated inflammatory response, particularly Interleukin (IL)-1Beta and IL-6. These latter have been shown, for instance, to be elevated in cancer patients ^24^, and administration of interferon-alpha produces depression and/or fatigue in the majority of patients receiving it as a treatment^25^. HPA axis dysregulation in the form of hypocortisolaemia, blunted diurnal variation and blunted stress reactivity have been found in cross-sectional studies of CFS patients (see Tomas et al.^26^ for a recent review). Other popular candidate aetio-pathological mechanisms for fatigue include serotonin pathways, circadian dysregulation, autonomic dysfunction^17^, 5HT neurotransmitter dysregulation, alterations in ATP metabolism, and vagal afferent activation^24^. Some authors have suggested that, rather than being located with one biological system, fatigue represents a systemic dysregulation of the interaction between these systems^27^.

At the other end of the biopsychosocial spectrum, psychosocial models of fatigue focus on the role that the individual’s response to their symptom may serve in perpetuating it. For instance, in a cross sectional study of 149 patients with multiple sclerosis, illness-related cognitions and behaviours were associated with a higher level of fatigue independent of neurological impairment^28^. More integrative models are predicated on the notion of allostatic load, the psychophysiological work done to adjust to stress, and its impact upon the body’s self-regulatory systems. As such, these models complement biological accounts of fatigue and provide potential pathways for integrating psychosocial and biological findings^29^. Multifactorial accounts and models of fatigue exist in multiple sclerosis^30^, primary biliary cirrhosis^14^, obesity^31^, diabetes^32^, frailty^33^ and cancer^13^. These multi-factorial models postulate that fatigue is likely to be the product of physiological factors (generic, such as inflammation, and/or disease specific such as hyperglycaemia in diabetes), psychosocial factors (e.g. emotional distress), lifestyle and behavioural factors (e.g. reduced activity), illness consequences (e.g. sleep disturbance, weakness), and the interaction of these contributors.

Research into the genetics and epigenetics of fatigue has tended to focus on genes associated with the biological mechanisms described above, and done so usually within fatiguing illnesses such as those listed above. Candidate gene studies have suggested several genes to be involved in chronic fatigue syndrome, particularly genes involved in the immune system and HPA axis. Reviewing this literature Landmark-Høyvik et al.^34^ suggest that findings are inconclusive, and are hampered by phenotypic heterogeneity, lack of power, and poor study design. Twin studies have shown heritability of fatigue to be between 6% and 50% with a higher concordance in monozygotic twins than dizygotic twins^35, 36^. In a study of fatigue, insomnia and depression in 3758 twins (893 MZ pairs and 884 DZ pairs) the best model was a common pathway model suggesting that the high association between the symptoms (correlations of 0.35 to 0.44) was mediated by an underlying common factor whose variation was 49% genetic and 51% unique environmental. Unique specific variance in fatigue was 38% genetic and 62% unique environmental^37^. Genome-wide association studies have shown an association between SNPs in genes associated with impaired cognitive abilities (*GRIK2, p* = 1.26 ×10^−11^)^38, 39^ and the circadian clock (*NPAS2*, not genome-wide significant)^39^ and chronic fatigue syndrome, but this was in a sample of just 42 cases of chronic fatigue syndrome and 38 controls.

To sum up in the words of Landmark-Høyvik et al.^40^: “fatigue can be conceptualized as a final common end point for psychological and biological processes. Fatigue is therefore both heterogeneous (occurring across different conditions) and multifactorial”. The aim of the present study is to understand further the genetic contribution to self-reported tiredness and/or low energy. We conducted a genome-wide association analysis, in the UK Biobank sample, of a response to a single item question: “Over the past two weeks, how often have you felt tired or had little energy?” Based on the foregoing literature overview, we also investigated pleiotropy with physical-and mental health-related traits.

## Materials and methods

### Study design and participants

UK Biobank is a large resource for identifying determinants of human diseases in middle aged and older individuals (http://www.ukbiobank.ac.uk)^41^. A total of 502 655 community-dwelling individuals aged between 37 and 73 years were recruited in the United Kingdom between 2006 and 2010. Baseline assessment included cognitive testing, personality self-report, and physical and mental health measures. For the present study, genome-wide genotyping data were available for 112 151 participants (58 914 females, 53 237 males) after quality control (see below). They were aged from 40 to 73 years (mean = 56.9 years, SD = 7.9). UK Biobank received ethical approval from the Research Ethics Committee (REC reference for UK Biobank is 11/NW/0382). This study has been completed under UK Biobank application 10279. Figure 1 shows the study flow for the present report.

**Figure 1.**
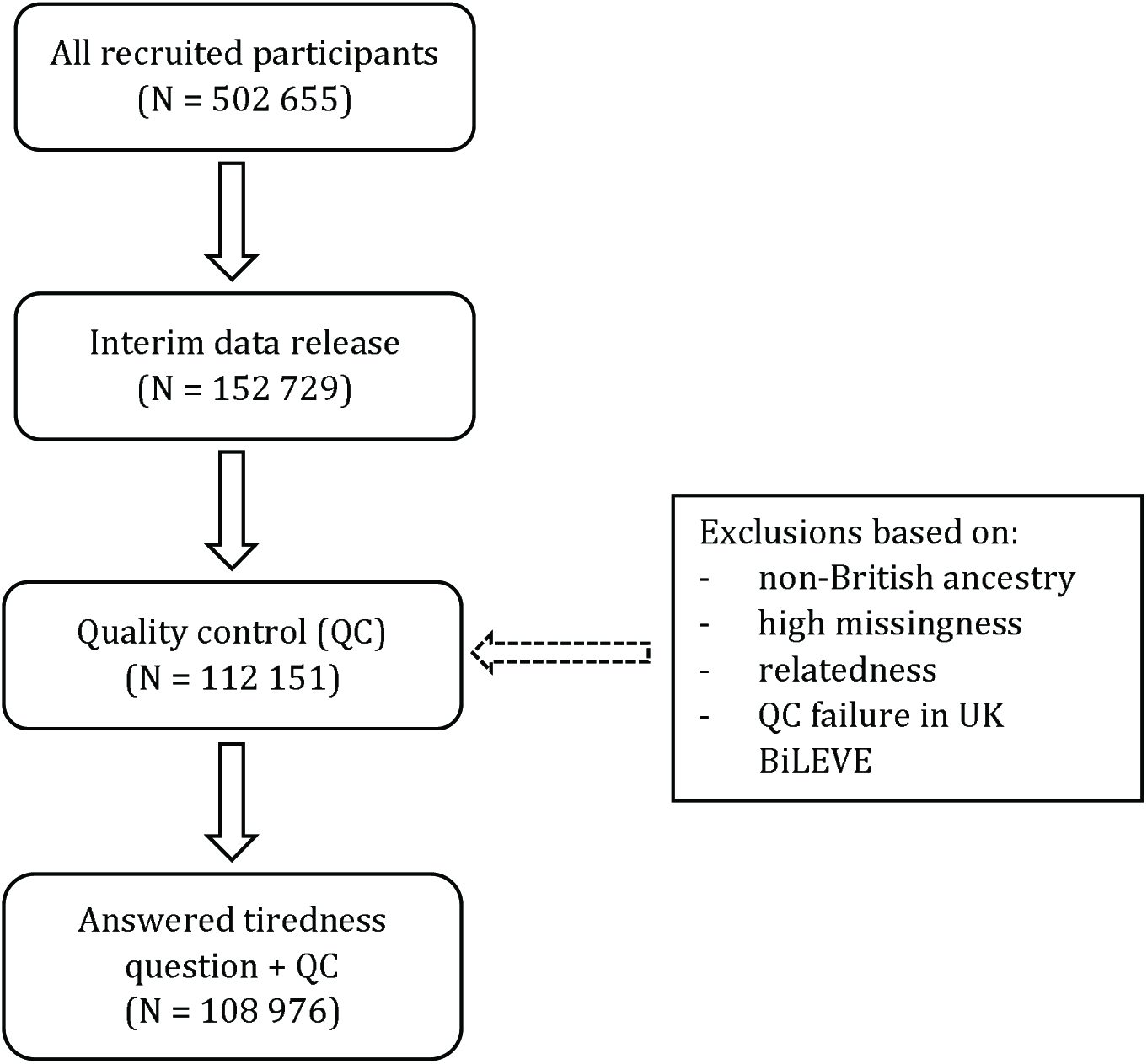
Flow diagram of participant selection.

### Procedures

#### Tiredness

Participants were asked the question, “Over the past two weeks, how often have you felt tired or had little energy?” Possible answers were: “Not at all/Several days/More than half the day/Nearly every day/Do not know/Prefer not to answer”. This question was asked as part of the Mental Health Questionnaire, which consists of items from the Patient Health Questionnaire^42^. Participants answering with “Do not know” or “Prefer not to answer” were excluded, resulting in a four-category variable for tiredness ranging from “Not at all” to “Nearly every day”. We will refer to this question in the rest of the paper as ‘tiredness’ but we ask the reader to bear in mind the question as it was asked, i.e. its referring to tiredness and/or low energy.

#### Genotyping and quality control

The interim release of UK Biobank included genotype data for 152 729 individuals, of whom 49 979 were genotyped using the UK BiLEVE array and 102 750 using the UK Biobank axiom array. These arrays have over 95% content in common. Details of the array design, genotyping procedures and quality control details have been published elsewhere ^43, 44^.

#### Imputation

An imputed dataset was made available as part of the UK Biobank interim data release. The 1000 Genomes phase 3 and UK10K haplotype reference panels were merged and the genotype data imputed to this merged reference panel. Further details can be found at the following URL: http://biobank.ctsu.ox.ac.uk/crystal/refer.cgi?id=157020. Autosomal variants with a minor allele frequency less than 0.1 % and an imputation quality score of 0.1 or less were excluded from further analysis (N ~ 17.3M SNPs).

#### Curation of summary results from GWAS consortia on health related variables

Published summary results from international GWAS consortia were gathered in order to perform linkage disequilibrium (LD) score regression and polygenic profile analysis between the UK Biobank tiredness variable and the genetic predisposition to multiple health-related traits. Details of the health-related variables, the consortia’s websites, key references for each consortium, and number of subjects included in each consortium’s GWAS are given in Supplementary Table 1.

### Statistical analysis

#### Phenotypic correlations

Spearman’s rank correlation coefficients were calculated between responses to the tiredness question, grip strength, forced expiratory volume in 1s, height, BMI, self-rated health, verbal-numerical reasoning and neuroticism, all of which were measured phenotypes in UK Biobank. Details on measurements of these phenotypes can be found in the Supplementary Material.

#### Genetic association analysis

111 749 individuals answered the tiredness question and had genotypic information. After visual inspection of the distribution of the UK Biobank tiredness variable no exclusions were made. Prior to analysis, tiredness was adjusted for age, gender, assessment centre, genotyping batch and array, and 10 principal components for population stratification. Genotype-phenotype association analyses were conducted using SNPTEST v2.5.1^45^ and can be found at the following URL: https://mathgen.stats.ox.ac.uk/genetics_software/snptest/snptest.html#introduction. An additive model was specified using the ‘frequentist 1’ option. Genotype uncertainty was accounted for by analysing genotype dosages.

Genetic association analyses were also performed on the following UK Biobank phenotypes in order to perform further analyses: self-rated health^46^, grip strength, forced expiratory volume in 1s, neuroticism^47^, verbal-numerical reasoning^48^. We specifically examined whether any variants associated with tiredness were also associated with grip strength, self-rated health, and neuroticism because we judged these to provide some coverage of physical and mental resilience in UK Biobank.

#### Estimation of SNP based heritability

To estimate the proportion of variance explained by all common SNPs in tiredness, univariate GCTA-GREML analysis was performed^49^. This analysis included only unrelated individuals, using a relatedness cut-off of 0.025 in the generation of the genetic relationship matrix.

#### Gene-based association analysis

Gene-based associations were derived using MAGMA^50^ using the summary GWAS statistics for tiredness. SNPs were assigned to 18 062 genes using the NCBI build 37.3. The gene boundary was defined as the start and stop site of each gene. In order to account for linkage disequilibrium between the SNPs used, the European panel of the 1000 Genomes data (phase 1, release 3) was used. A Bonferroni correction was used to control for 18 062 tests (α= 0.05/18,062; P < 2.768 x 10-6).

#### Partitioned heritability

The summary statistics from the GWAS on tiredness were portioned into functional categories using the same data processing pipeline as Finucane et al.^51^; more details on this method can be found in the Supplementary Materials.

#### Pleiotropy; LD score regression and polygenic profiling

Genetic associations between tiredness and health-related variables from GWAS consortia were computed using two methods, LD score regression and polygenic profile score analysis. Each provides a different metric to infer the existence of pleiotropy between pairs of traits. LD score regression was used to compute genetic correlations between two traits; this tests the degree to which the polygenic architecture of one trait overlaps with that of other traits. Polygenic profile score analysis was used to test the extent to which individual differences in the tiredness phenotype in UK Biobank could be predicted by the polygenic architecture of the health-related traits from other GWAS consortia. Both of these methods are dependent on a trait’s being highly polygenic in nature, that is, where a large number of variants of small effect contribute toward phenotypic variation^52, 53^. LD score regression was performed between tiredness and 28 health-related traits. Polygenic profile score analysis was performed on 25 of the 28 health-related traits, as this method requires independent samples to provide the summary GWAS information from which the polygenic profile score is based.

##### LD score regression

This was used to quantify the extent of pleiotropy between tiredness in UK Biobank and 28 health-related traits^53, 54^. This technique examines the correlational structure of the SNPs found across the genome. In the present study, LD score regression was used to derive genetic correlations between tiredness and health-related traits using the GWAS results of 24 large GWAS consortia and four UK Biobank phenotypes. The data processing pipeline devised by Bulik-Sullivan et al.^53^ was followed. In order to ensure that the genetic correlation for the Alzheimer’s disease phenotype was not driven by a single locus or biased the fit of the regression model, a 500kb region centred on the *APOE* locus was removed and this phenotype was re-run. This additional model is referred to in the tables and figures as ‘Alzheimer’s disease (500kb)’.

##### Polygenic profile scores

The UK Biobank genotyping data was recoded from numeric (1,2) allele coding to standard ACGT coding using a bespoke programme developed by one of the present authors (DCL)^44^. Polygenic profile scores were created for 25 health-related traits in all genotyped participants using PRSice^55^. Prior to creating the scores, SNPs with a minor allele frequency < 0.01 were removed and clumping was used to obtain SNPs in linkage equilibrium with an r^2^ < 0.25 within a 200bp window. Five polygenic profile scores were created for each trait including SNPs according to their significance of association with the relevant trait at p-value thresholds of P < 0.01, P < 0.05, P < 0.1, P < 0.5 and all SNPs.

Regression models were used to examine the association between the polygenic profiles and tiredness in UK Biobank, adjusting for age at measurement, sex, genotyping batch and array, assessment centre, and the first ten principal components for population stratification. All associations were corrected for multiple testing using the false discovery rate method^56^. Sensitivity analyses were performed to test whether the results were confounded by individual’s neuroticism levels, their self-rated health scores, or a diagnosis of major depressive disorder. This was done by adjusting the models for the neuroticism and self-rated health scores. Individuals with a probable diagnosis of major depressive disorder were excluded from the sensitivity analysis, based on the diagnostic method formulated by Smith et al.^57^. Further details can be found in the Supplementary Material. To examine whether any association between polygenic risk for type 2 diabetes and tiredness was confounded by having had a diagnosis of type 2 diabetes, all individuals with a self-reported doctor’s diagnosis of type 2 diabetes were excluded from that specific sensitivity analysis (Supplementary Material). Multivariate regression was performed using all FDR significant polygenic profile scores and earlier described covariates.

#### Comparison of gene-based analysis results within UK Biobank

Gene-based associations for tiredness were compared with gene-based results for other UK Biobank health-related traits that, in the present report’s results, showed a statistically significant genetic correlation with tiredness, using previously described methods^50^.

## Results

### Phenotypic correlations

108 976 individuals from UK Biobank with genotypic data answered the question “Over the past two weeks, how often have you felt tired or had little energy?”, referred to hereinafter as “tiredness”. There were 51 416 individuals who answered “not at all”, 44 208 individuals responded “several days”, 6404 individuals answered “more than half the days”, and 6948 individuals responded “nearly every day”. Correlations indicated that individuals who reported feeling more tired tended to have lower grip strength, lower lung function, poorer self-rated health, lower scores for verbal-numerical reasoning, and shorter stature (Table 1). Correlations indicated that individuals who reported feeling more tired tended to have a higher BMI and higher neuroticism scores (Table 1). Absolute effect sizes ranged from very small to moderate (Supplementary Figures 1a-f). The mean scores and distribution for each of these variables at each level of tiredness are shown in Supplementary Figure 1 and Supplementary Table 2.

**Table 1.** Spearman phenotypic correlations between tiredness (responses to the question, “Over the past two weeks, how often have you felt tired or had little energy?”) and physical and mental health. All correlations had P < 0.001

|  | **Tiredness** |
| --- | --- |
| **Self-rated health (N = 108 648)** | -0.35 |
| **Grip strength (N = 108 573)** | -0.12 |
| **Forced expiratory function in 1s (N = 101 823)** | -0.08 |
| **Height (N = 108 796)** | -0.09 |
| **BMI (N = 108 681)** | 0.10 |
| **Verbal-numerical reasoning (N = 35 101)** | -0.04 |
| **Neuroticism (N = 105 456)** | 0.39 |

### Genome-wide association study

There was one genome-wide significant SNP (Affymetrix id 1:64178756_C_T; p = 1.36 x 10^−11^) on chromosome 1 (Figure 2). This SNP is not in a gene and does not have an rs id. It has both a low minor allele frequency (0.001) and a low imputation quality score (0.43). It is not in a peak with other SNPs. Therefore, this result should be treated with caution. Two suggestive peaks were identified on chromosomes 1 and 17, with the lowest p values of 5.88 x 10^−8^ (rs142592148; an intronic SNP in S*LC44A5*) and 6.86 x 10^−8^ (rs7219015; an intronic SNP in *PAFAH1B1*) for each peak, respectively. The peak on chromosome 1 contains three genes (*CRYZ, TYW3 and SLC44A5*) and the peak on chromosome 17 contains one gene (*PAFAH1B1*). The *CRY/TYW3* locus has previously been associated with circulating resistin levels, a hormone associated with insulin resistance, inflammation, and risk of type 2 diabetes and cardiovascular disease^58^. *SLC44A5* encodes a solute carrier protein and is important for metabolism of lipids and lipoproteins and has been associated with birth weight in cattle ^59^. *PAFAH1B1* encodes a subunit of an enzyme that has important roles in brain development and spermatogenesis. Mutations in this gene cause the neurological disorder lissencephaly^60^.

**Figure 2.**
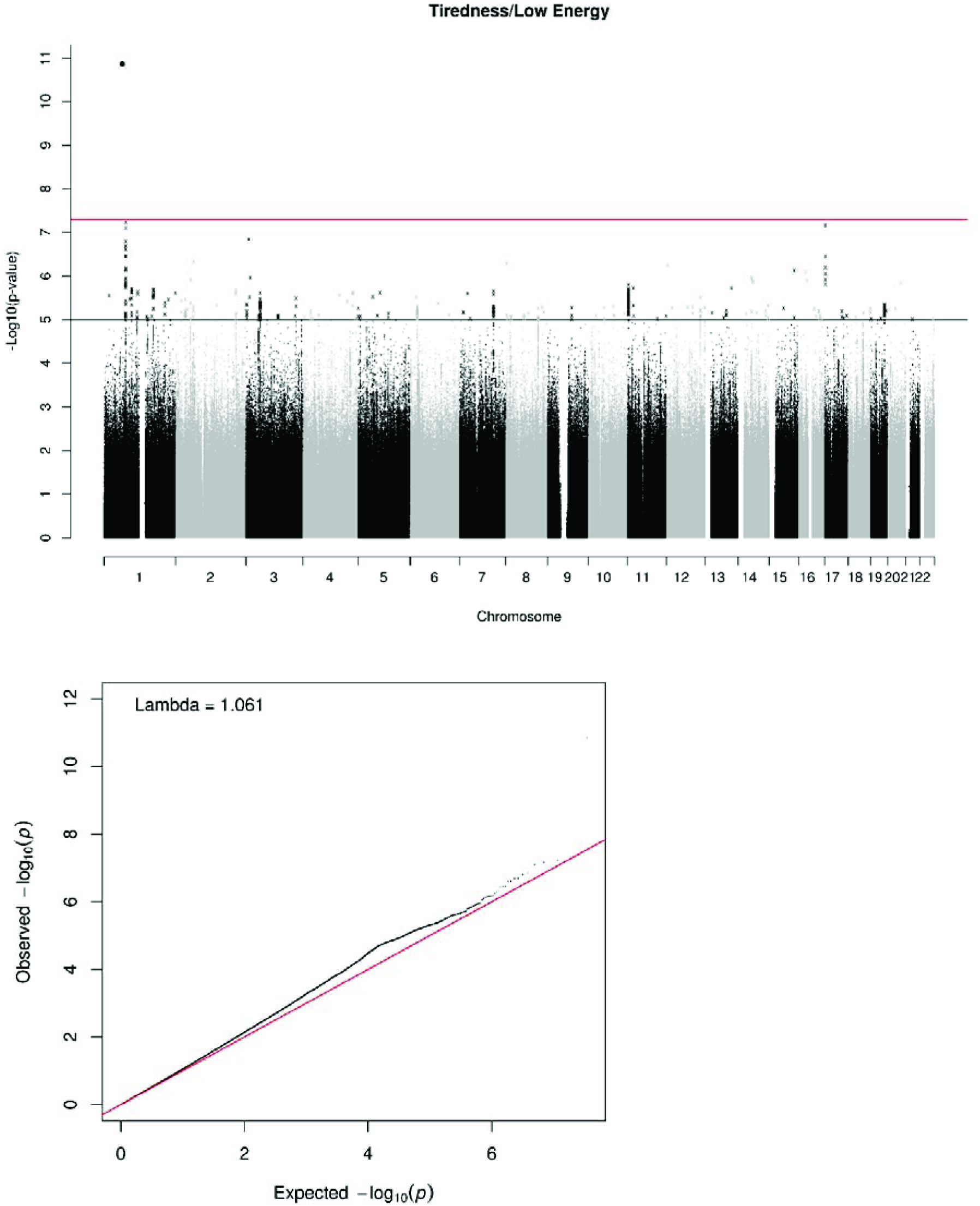
(A) Manhattan and (B) Q-Q plot of P-values of the SNP-based association analysis of tiredness (responses to the question, “Over the past two weeks, how often have you felt tired or had little energy?”). The red line indicates the threshold for genome-wide significance (P<5 x 10-8); the grey line indicates the threshold for suggestive significance (P<1 x 10-5).

### SNP based heritability estimate

The proportion of variance in tiredness explained by all common genetic variants using GCTA-GREML was 8.4 % (SE 0.006).

### Gene based association analysis

Gene-based association analysis identified five genes, *DRD2*, *PRRC2C*, *C3orf84*, *ANO10* and *ASXL3*, that attained genome-wide significance for tiredness (Table 2 and Supplementary Table 3). *DRD2* encodes a dopamine receptor and has previously been associated with psychiatric illnesses^61^. Alternative splicing of *PRRC2C* has been associated with lung cancer^62^. Mutations in *ANO10* cause cerebellar ataxias^63^. *ASXL3* encodes a polycomb protein and mutations in this gene are associated with intellectual disability, feeding problems and distinctive facial features^64^.

**Table 2.** Shows the genome wide significant genes from the UK Biobank tiredness phenotype and the significance values for the same genes using the neuroticism, self-rated health, and grip strength phenotypes, also in the UK Biobank sample. Unmodified P-values are shown for all phenotypes.

| CHR | Gene | Tiredness P | SRH P | Grip P | Neuroticism P |
| --- | --- | --- | --- | --- | --- |
| 11 | *DRD2* | $2.94 \times 10^{-7}$ | 0.012 | 0.156 | $9.69 \times 10^{-9}$ |
| 1 | *PRRC2C* | $1.43 \times 10^{-6}$ | 0.002 | 0.314 | 0.020 |
| 3 | *C3orf84* | $1.45 \times 10^{-6}$ | $7.38 \times 10^{-5}$ | 0.001 | 0.016 |
| 3 | *ANO10* | $1.52 \times 10^{-6}$ | 0.058 | 0.728 | 0.001 |
| 18 | *ASXL3* | $2.67 \times 10^{-6}$ | $1.03^{-6}$ | 0.052 | $1.36 \times 10^{-5}$ |

In addition, each of these genes was also nominally significant in a GWAS of neuroticism, with *DRD2* being genome wide significant in both phenotypes^47^. A comparison with a UK Biobank GWAS of self-rated health^22^ showed that four of the five genes (*DRD2*, *PRRC2C*, *C3orf84* and *ASXL3*) were significant across phenotypes. Grip strength showed less overlap, with only *C3orf84* being nominally associated with grip strength. Of the five genes examined here, *C3orf84* was associated with each of the four phenotypes. Whereas the genetic correlations between these traits (see below) are likely to encompass multiple genic and non-genic regions, as well as unique points of overlap between pairs of phenotypes, the variants found in the *C3orf84* represent a point of the genome where the genetic architecture of these four traits converges.

### Partitioned heritability

From the full baseline model using 52 annotations, only evolutionarily conserved regions were found to be enriched for tiredness (Supplementary Figure 2). This annotation contained only 2.6% of the SNPs from the summary statistics, but they collectively explained 40% of the heritability of tiredness (SE = 11%, enrichment metric = 15.34, SE = 4.05, P = 0.00039). By clustering the histone marks into tissue specific categories we found significant enrichment for variants found in the central nervous system (Supplementary Figure 3). This category contained 15% of the SNPs and explained 45% of the heritability (SE = 8%, enrichment metric = 3.02, SE = 0.54, P = 0.0002).

### Genetic correlations between tiredness and physical and mental health traits

LD score regression was used to test if genetic variants associated with health-related traits also contribute towards tiredness in UK Biobank. Table 3 and Figure 3 show these genetic correlations. Positive significant (FDR corrected) genetic correlations were found between tiredness and BMI (r_g_ = 0.20), HbA1c (r_g_ = 0.25), obesity (r_g_ = 0.21), smoking status (r_g_ = 0.20), triglycerides (r_g_ = 0.13), type 2 diabetes (r_g_ = 0.18), waist-hip ratio (r_g_ = 0.08), ADHD (r_g_ = 0.27), bipolar disorder (r_g_ = 0.14), major depressive disorder (r_g_ = 0.59), neuroticism (r_g_ = 0.62), and schizophrenia (r_g_ = 0.25). Negative significant (FDR corrected) genetic correlations were found between HDL cholesterol (r_g_ = −0.11), forced expiratory volume in 1s (r_g_ = −0.12), grip strength (r_g_ = −0.16), longevity (r_g_ = −0.39), self-rated health (r_g_ = −0.78), and verbal-numerical reasoning (r_g_ = 0.14). These genetic correlations suggest that there are common genetic associations between tiredness and multiple physical-and mental health-related traits.

**Figure 3.**
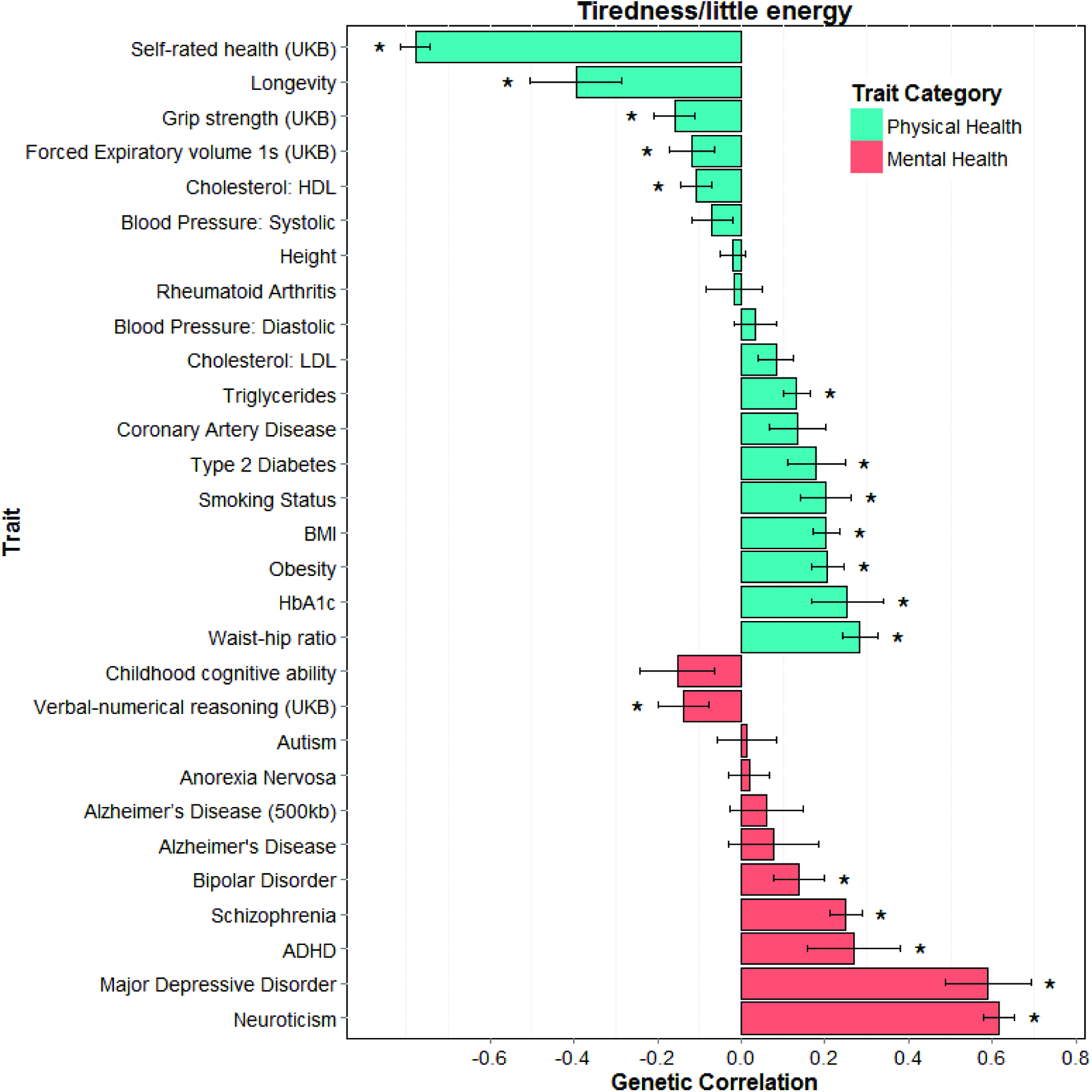
Barplot of genetic correlations (SE) calculated using LD regression between tiredness in UK Biobank and mental and physical health measures from GWAS consortia. *, p< 0.0281

**Table 3.**
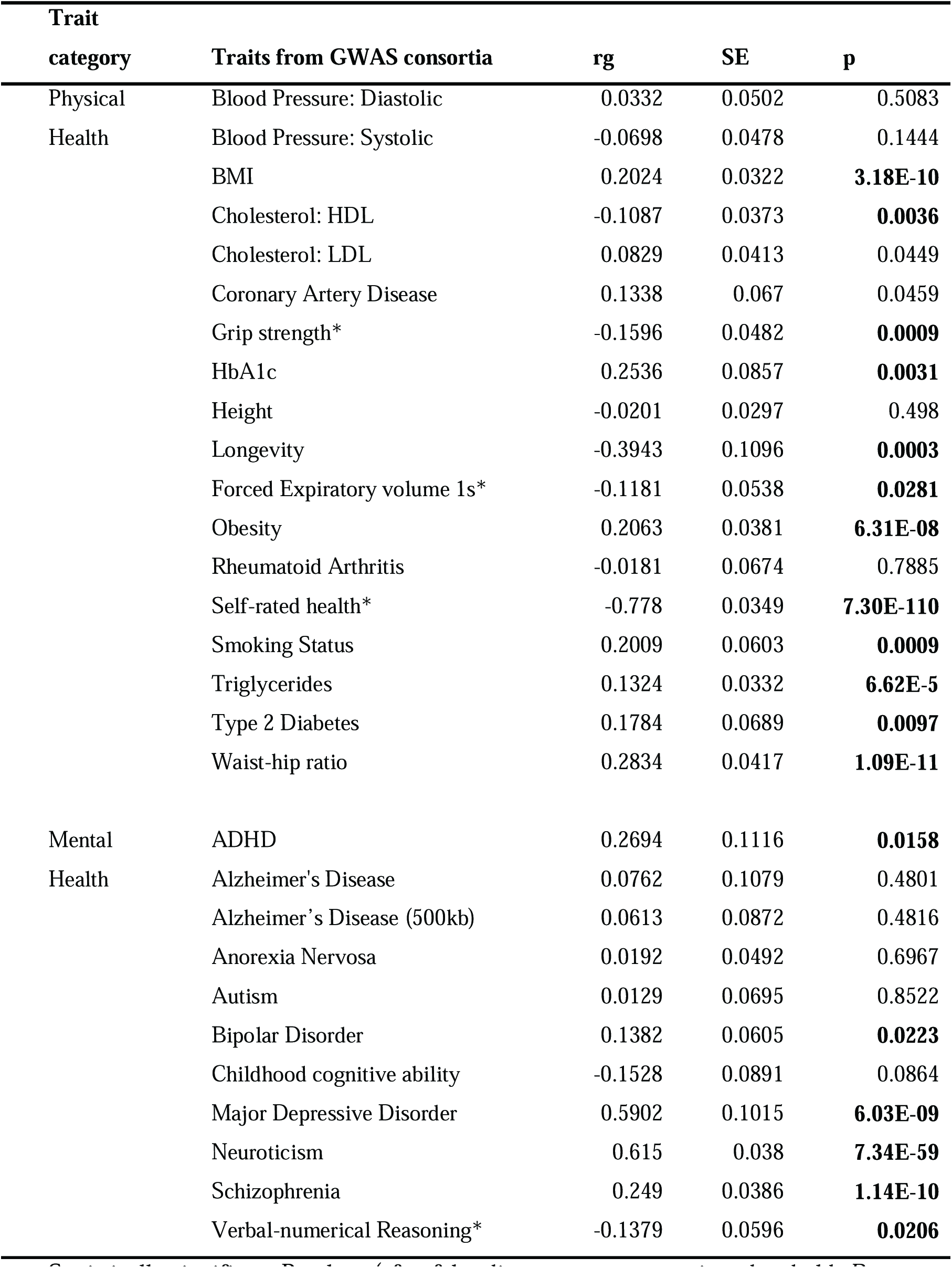
Genetic correlations between tiredness documented in the UK Biobank data set and the health-related variables collected from GWAS consortia.

Statistically significant P-values (after false discovery rate correction; threshold: P = 0.0281) are shown in bold. *GWAS based on UK Biobank data.

### Polygenic prediction

The full results including all five thresholds can be found in Supplementary Table 4, as well as the number of SNPs included for the five thresholds in each trait. Table 4 shows the results for the polygenic profile scores analyses, using the most predictive threshold for each trait. Complete results for all five thresholds can be found in Supplementary Table 4. Higher polygenic profiles for nine physical health traits predicted increased tiredness (standardised β’s between 0.008 and 0.03) in UK Biobank: BMI, LDL cholesterol, coronary artery disease, HbA1c, obesity, smoking status, triglycerides, type 2 diabetes, and waist-hip ratio. Higher polygenic profiles for HDL cholesterol and height predicted lower tiredness (β = −0.016 and β = −0.008, respectively).

**Table 4.** Associations between polygenic profiles of health-related traits created from GWAS consortia summary data, and the UK Biobank tiredness phenotype controlling for age, sex, assessment centre, genotyping batch and array and 10 principal components for population structure

| Trait Category | Trait | Threshold | $\beta$ | P |
| --- | --- | --- | --- | --- |
| Physical Health | Blood Pressure: Diastolic | 0.1 | -0.0028 | 0.3619 |
|  | Blood Pressure: Systolic | 0.1 | -0.0025 | 0.4077 |
|  | BMI | 1 | 0.0280 | **4.90×10<sup>-20</sup>†** |
|  | Cholesterol: HDL | 0.5 | -0.01634 | **8.49×10<sup>-8</sup>†** |
|  | Cholesterol: LDL | 0.5 | 0.008147 | **0.0077<sup>†</sup>** |
|  | Coronary Artery Disease | 0.5 | 0.0084 | **0.0061** |
|  | Forced Expiratory volume 1s | 0.01 | -0.0059 | 0.0529 |
|  | Longevity | 0.05 | -0.0067 | 0.0297 |
|  | HbA1c | 1 | 0.008999 | **0.0033<sup>†</sup>** |
|  | Height | 1 | -0.0077 | **0.0154** |
|  | Obesity | 1 | 0.0236 | **1.20×10<sup>-14</sup>†** |
|  | Rheumatoid Arthritis | 0.1 | -0.0016 | 0.5926 |
|  | Smoking Status | 0.5 | 0.0091 | **0.0048** |
|  | Triglycerides | 0.5 | 0.020854 | **1.06×10<sup>-11</sup>†** |
|  | Type 2 Diabetes | 1 | 0.0120 | **0.0001<sup>†a</sup>** |
|  | Waist-hip ratio | 1 | 0.025791 | **7.85×10<sup>-17</sup>** |
| Mental Health | ADHD | 1 | 0.0042 | 0.1647 |
|  | Alzheimer's Disease | 0.05 | -0.0052 | 0.0889 |
|  | Anorexia Nervosa | 0.5 | 0.0048 | 0.1169 |
|  | Autism | 1 | -0.0018 | 0.5593 |
|  | Bipolar Disorder | 0.01 | 0.0082 | **0.0076\*** |
|  | Childhood cognitive ability | 0.1 | -0.0112 | **0.0002<sup>†</sup>** |
|  | Major Depressive Disorder | 1 | 0.0183 | **2.00×10<sup>-9</sup>\*** |
|  | Neuroticism | 0.1 | 0.0185 | **2.25×10<sup>-9</sup>\*** |
|  | Schizophrenia | 1 | 0.0290 | **3.25×10<sup>-20</sup>†\*** |

FDR-corrected statistically significant values (P=0.0255) are shown in bold. The associations between the polygenic profile with the largest effect size (threshold) and tiredness are presented. Threshold is the P-value threshold with the largest effect size. †, results remain significant after controlling for neuroticism scores; *, results remain significant after controlling for self-rated health; ^a^, results remain significant after excluding individuals with type 2 diabetes (β = 0.0105, p = 0.00076).

Of the mental health traits, higher polygenic risk for bipolar disorder, neuroticism, major depressive disorder, and schizophrenia were associated with increased tiredness (standardised β’s between 0.008 and 0.03). Polygenic profiles for childhood cognitive ability showed a negative association with tiredness (β = −0.01).

Sensitivity analysis showed that, when controlling for neuroticism, the associations between tiredness and polygenic profiles for BMI, obesity, type 2 diabetes, cholesterol (HDL and LDL), HbA1c, triglycerides, waist-hip ratio, childhood cognitive ability, and schizophrenia remained significant (after FDR correction), indicating that these associations are not confounded by scores for neuroticism. Similar analyses controlling for self-rated health indicated that the following associations with tiredness are not confounded by self-rated health: bipolar disorder, neuroticism, major depressive disorder, and schizophrenia. Supplementary Table 5 shows the adjusted results and the percentage of attenuation in standardised β’s for the models.

When excluding individuals with a probable diagnosis of major depressive disorder (N = 7364) from all individuals with sufficient information about their mental health to make a probable depression diagnosis (N = 31 523), the associations between tiredness and polygenic profiles for BMI, HDL cholesterol, obesity, triglycerides, waist-hip ratio, neuroticism, and schizophrenia remained significant (after FDR correction), compared to BMI, HDL cholesterol, obesity, triglycerides, type 2 diabetes, waist-hip ratio, neuroticism, and schizophrenia in the full model (also N = 31 523) (Supplementary Table 6). When excluding individuals with a type 2 diabetes diagnosis (N = 725), the association between tiredness and the polygenic risk score for diabetes remained significant, indicating that this association is independent of self-reported morbidity for that disorder.

A multivariate regression model including 15 of the 16 significant polygenic profile scores (BMI, HDL cholesterol, LDL cholesterol, HbA1c, height, obesity, smoking status, triglycerides, type 2 diabetes, waist-hip ratio, bipolar disorder, childhood cognitive ability, major depressive disorder, neuroticism and schizophrenia) showed that polygenic profile scores for the following traits contributed independently to the association with tiredness: BMI, HDL cholesterol, triglycerides, waist-hip ratio, childhood cognitive ability, major depressive disorder, neuroticism, and schizophrenia. The scores together accounted for 0.25 % of the variance in tiredness (Supplementary Table 7).

## Discussion

In the present study, pleiotropy was identified between tiredness and longevity, grip strength, multiple metabolic indicators, smoking status, neuroticism, childhood cognitive ability, depression, and schizophrenia. These analyses, combining data from the UK Biobank and many GWAS consortia, provide the first estimate of the overlap in the genetic variants contributing to the heritability of tiredness and these physical and mental health-related traits and disorders. Tiredness demonstrated significant SNP-based heritability of 8.4 %.

Five genes attained genome wide significance for tiredness: *DRD2*, *PRRC2C*, *ANO10*, *ASXL3* and *C3orf84*, the latter being an uncharacterised protein representing a point of genetic convergence between tiredness, neuroticism, grip strength and self-rated health. *DRD2*, *PRRC2C*, *ANO10*, *ASXL3*, and *DRD2* have previously been associated with psychiatric illnesses^18^, lung cancer^19^, cerebellar ataxias^20^ and intellectual disability^21^ respectively. The three genes within the suggestive peak on chromosome 1 (*CRYZ, TYW3* and *SLC44A5*) have previously been associated with insulin resistance, inflammation, risk of type 2 diabetes, metabolism of lipids and lipoproteins and cardiovascular disease^15,16^. These genes are consistent with the identification of pleiotropy between tiredness and metabolic irregularities, and perhaps more broadly with ‘metabolic syndrome’ and ‘allostatic load’. *PAFAH1B*, within a suggestive peak on chromosome 17 has important roles in brain development^17^. This is consistent with the identification of pleiotropy between tiredness and cognitive traits and the finding of significant enrichment for variants found in the central nervous system.

Evolutionarily conserved regions were found to be enriched for association with tiredness, consistent with findings for other quantitative traits including disease status^51^, suggesting that these are important loci where mutations result in detrimental phenotypes. The range of factors—affective, cognitive, behavioural, and physical—that are genetically associated with tiredness is in itself remarkable, and confirms the observation of Landmark-Høyvik et al^34^, quoted in the introduction, that the related construct of fatigue is aetiologically heterogeneous and multifactorial. No overlap has been found with genes (*GRIK2*, *NPAS2*) identified in previous GWAS of fatigue ^38, 39^; however, the sample size of these studies was too small to have enough power to detect statistically significant differences. The present study did not show significant associations for candidate genes previously identified^34^.

Tiredness showed significant shared heritability with a range of factors associated with metabolic syndrome^65^ including cholesterol, tri-glycerides, HbA1c, waist-hip ratio, BMI, obesity, and type 2 diabetes. Several of these factors are biomarkers of allostatic load ^61, 62^. The concept of allostatic load has been used in the context of both physical and mental ill health, including the symptom of fatigue. Conceptually, allostatic load represents the cumulative, physiological ‘wear and tear’ of a prolonged response to a stressor. Allostatic load has been shown to be a replicably-measurable multi-variate construct^66, 67^, with first order factors comprising cardiovascular, immune, metabolic, anthropometric, and neuroendocrine markers^29^. It is hypothesised that, in response to threats to homeostasis, the body’s self-regulatory mechanisms, such as the sympathetic-adrenal medullary axis and the hypothalamic pituitary adrenal axis, have the potential to “overcompensate and eventually collapse upon themselves”^29^, with consequences for morbidity and mortality.

In our current analysis we included metabolic (cholesterol, HbA1c, triglycerides), anthropometric (waist hip ratio, BMI, obesity) and cardiovascular/respiratory (diastolic and systolic blood pressure, forced expiratory volume) markers of allostatic load. The results showed significant pleiotropy, as measured by both LD score regression and polygenic profile analysis, between tiredness and most of the metabolic and anthropometric markers, though not the cardiovascular/respiratory markers. This is suggestive of the fact that the pleiotropy between tiredness and these physiological factors may be due to a biological propensity to an over-compensatory physiological stress response.

Neuroticism, the tendency to experience negative affective states, and tiredness were strongly correlated, both phenotypically and genetically. This may represent a separate route to fatigue, a predominately affective one, and/or it may overlap with the concept of allostatic load. A recent paper by Gale et al.^47^, also using this UK Biobank sample, supports the allostatic load link with neuroticism, showed that polygenic profile scores for several physical and mental health traits - BMI, coronary artery disease, smoking status, bipolar disorder, borderline personality, major depressive disorder, negative affect and schizophrenia - significantly predicted neuroticism. In the present study, when tiredness polygenic profile analyses were adjusted for neuroticism, the associations between tiredness and mental health disorders (bar schizophrenia) were largely attenuated, whereas most of the metabolic and anthropometric associations remained significant. This suggests that it is the propensity to neuroticism, rather than the specific propensity to these disorders, that accounts or mediates the tiredness associated with mood disorders. Watson and Pennebaker^68^, discussing competing models of how negative affectivity is related to self-reported physical and emotional well-being, found that it is associated as much with the former as with the latter, and that negative affectivity might better be conceptualised as a general tendency to experience both somatic and emotional distress. This concept of a general tendency to what they termed somatopsychic distress could explain the pleiotropy observed in the present study between neuroticism and tiredness.

That neuroticism may be a distinct route to fatigue is supported by the fact that when the polygenic profile analysis is adjusted for self-rated health, all associations between polygenic profiles for physical health and tiredness are attenuated to the point of non-significance, whereas the relationship between tiredness, and polygenic profiles for neuroticism, MDD and bipolar disorder remain significant. This is consistent with the study of Gale et al.^69^, investigating pleiotropy between neuroticism and physical and mental health, where there were more and stronger genetic associations between neuroticism and mental health than between neuroticism and physical health. If we take self-rated health to be a marker, to some extent, of actual physical health (and the study by Harris et al.^70^ would indicate that it is), then this would suggest that when physical health is adjusted for, polygenic profiles for neuroticism and its associated negative affective states, continue to make a unique contribution to tiredness.

These proposed affective and physiological routes to fatigue may not be mutually exclusive. The allostatic load concept postulates that individual differences in personality, cognition and behavioural responses to stress, and socio-cultural factors, affect the physiological stress response. An increased propensity to experience distress, as captured in the concept of neuroticism, would imply that there is increasing propensity to over-respond to stressors and thus to physiological dysregulation. In the study by Gale et al.^69^, the only significant pleiotropy between neuroticism and physical disease state was a significant association with the polygenic risk score for coronary artery disease, which is suggestive of an overlap of affective and cardiovascular stress responses. The allostatic load model also allows us to incorporate childhood cognitive ability (reduced ability to problem solve), smoking status (at least one study has found smoking and allostatic load interaction effects^29^) and grip strength (reduced overall system integrity/vigour) into a more general model which is suggestive of shared genetic variance between stress proneness (neuroticism, reduced cognitive ability, reduced vigour), the physiological response to stress (biomarkers), behavioural responses (smoking), self-reported tiredness, disease and mortality^71^.

The large sample size of the present study is a strength of this study, providing powerful and robust tests of pleiotropy between tiredness and physical and mental health. A second strength is that all genetic samples were processed on the same platform at the same location. The use of summary data from many international GWAS consortia provided foundations for a comprehensive examination of shared genetic aetiology between tiredness and a wide range of health-related phenotypes.

A limitation of the present study is the measure of tiredness. As Wessely^72^ observed, an objective measure of fatigue is “an unattainable holy grail”. Almost all the studies cited in the introduction have used subjective self-reports. The self-report measures used vary widely, with there being several validated fatigue measures, and many of the reported studies use either double or single item questionnaires and/or single item Visual Analogue Scales. This in itself may account for some of the inconsistency in fatigue research, though the demographic studies cited at the beginning of this article, using a wide variety of measures from single questions^7^ to a well validated fatigue questionnaire^2^, produced similar findings. However our findings of genetic associations of the “wabbit” proxy for fatigue will need replication with better validated multi-item measures. All analyses were restricted to individuals of white British ancestry, because the sample does not have enough power to generalize results for individuals with different backgrounds.

## Summary

Being genetically predisposed to a range of mental and physical health complaints also predisposes individuals to report that they are more tired or lacking in energy. This study confirms that self-reported tiredness is a partly-heritable, heterogeneous and complex phenomenon that is phenotypically and genetically associated with affective, cognitive, personality, health and physiological processes. This study also served as a first step in testing some genetic hypotheses from the allostatic load model, finding suggestive links between tiredness and three genes on chromosome one associated with allostatic processes and considerable pleiotropy between tiredness and allostatic markers. We can foresee more tests of these links as more genome-wide genotyping data become available.

## Acknowledgments

This research has been conducted using the UK Biobank Resource. The work was undertaken in The University of Edinburgh Centre for Cognitive Ageing and Cognitive Epidemiology, part of the cross council Lifelong Health and Wellbeing Initiative (MR/K026992/1). Funding from the BBSRC, Age UK (Disconnected Mind Project), and Medical Research Council (MRC) is gratefully acknowledged.

